# Proteome remodelling in *Candida auris* during early host adaptation *in-vitro* and *in-vivo*

**DOI:** 10.64898/2026.03.09.710370

**Authors:** Rounik Mazumdar, Ana Bjelanovic

**Author notes:** **Correspondence:** Rounik Mazumdar.

## Abstract

*Candida auris* is an emerging fungal pathogen posing a serious global health threat due to its high transmissibility and multidrug resistance profile. Despite recent molecular advances in scrutinizing this enigmatic microbe, much of our understanding in regards to its pathomechanisms still remain unelucidated. Since, microbial pathogenesis is modulated by a dynamic interplay between the host and the pathogen, dissecting such host-pathogen interaction involving *C. auris* can shed novel insights into its pathogenic cascade. As such, to further characterize the virulence repertoire of *C. auris*, this study applied an integrated quantitative proteomics strategy to scrutinize early-phase of infection. We utilized an *in-vitro* and an *in-vivo* experimental setup based on immune cells and murine model. Integrated proteomic analysis revealed a coordinated remodelling of cellular processes by *C. auris* during host-pathogen interaction, including downregulation of translational machinery, and modulation of molecules involved in metabolic rewiring, stress-response, and structural rearrangements. Collectively, these findings suggests that survival of *C. auris* under host-immune pressure is accompanied by rapid context-dependent molecular adaptations.

**Importance:** *Candida auris* is a critical high priority fungal pathogen classified by the World Health Organization (WHO) that constitute a serious threat to global health. Often termed as a ‘superbug’ due to its high transmissibility and multidrug resistant profile, the microbe has spread across the globe and is capable of causing high mortality rates. Molecular studies scrutinizing the pathogenic mechanisms of *C. auris* are limited and represents a major bottleneck to decipher and device intervention strategies against this enigmatic pathogen. As such, this study is aimed at widening the molecular knowledge spectrum of *C. auris* in regards to its virulence and pathogenesis. Here we dissect the host-pathogen interaction of *C. auris* by establishing experimental infection models and subsequently applying an integrated proteomics strategy to capture the organism’s virulence repertoire modulating fungal pathogenesis.

## 1 Introduction

*Canida auris (Candidozyma auris)* a fungal pathogen first identified in 2009 in Japan, has ever since emerged as a ‘superbug’ proliferating across the globe and causing severe nosocomial outbreaks [1]. Genomic analysis of *C. auris* clinical strains have identified six clades, based on geographic locations including clade I (South Asia), clade II (East Asia), clade III (Africa), clade IV (South America), clade V (Iran) and the very recent clade VI (Singapore), with clade I being the most prevalent [2–4]. The pathogen can cause deep-seated infections of the bloodstream (candidemia), wound, respiratory tract, urinary tract, and ear infections, with high mortality rates ranging from 28% to 56% [5,6]. The fungi have been listed by the World Health Organization (WHO) as a ‘critical priority’ pathogen in its fungal priority pathogens list (FPPL), warranting urgent attention and research. Despite recent advances in our understanding of pathomechanisms of *C. auris*, much still remails unelucidated. The pathogen is particularly problematic owing to its high transmissibility, clinical misdiagnosis and drug resistance profile, often displaying multidrug resistance (MDR). It has been reported that 90% of *C. auris* clinical isolates are resistant to fluconazole, 35% to Amphotericin B (AmB), 7% to echinocandins, 3% to flucytosine and over 40% were reported to be resistance to more than two classes of antifungals [7]. Treatment options can therefore be limited by intrinsic or secondary resistance and due to the availability of low number of antifungal families. Understandably the current trends of research have been oriented towards deciphering the drug resistance mechanism of *C. auris* [8,9]. However, to effectively combat the pathogen a rigorous understanding of its virulence cascade is also warranted. Emerging reports are pointing towards a very complex pathogenic cascade harboured by the microbe [10]. Evidence suggests *C. auris* can efficiently adapt to host-microenvironment in order to evade host-immune system, establish and maintain an infection [10]. Henceforth, in addition to the ongoing focus on addressing drug resistance of *C. auris*, we also need to tackle the virulence aspect of the pathogen. As such, this study focuses on the molecular basis of virulence in *C. auris*, aiming to shed novel insights into its pathogenic cascade. To address this, we applied a quantitative proteomics strategy to identify the virulence factors modulating host-pathogen interaction during early-phase of infection. We established infection models using *in-vitro* co-culture setups with immune cells and an *in-vivo* murine infection model (Figure 1). Following a brief exposure of *C. auris* to host cells both *in-vitro* and *in-vivo*, we subjected reisolated fungal cells to quantitative proteomics analysis to identify molecules mediating host-pathogen interaction during the initial stages of infection.

**Figure 1.**
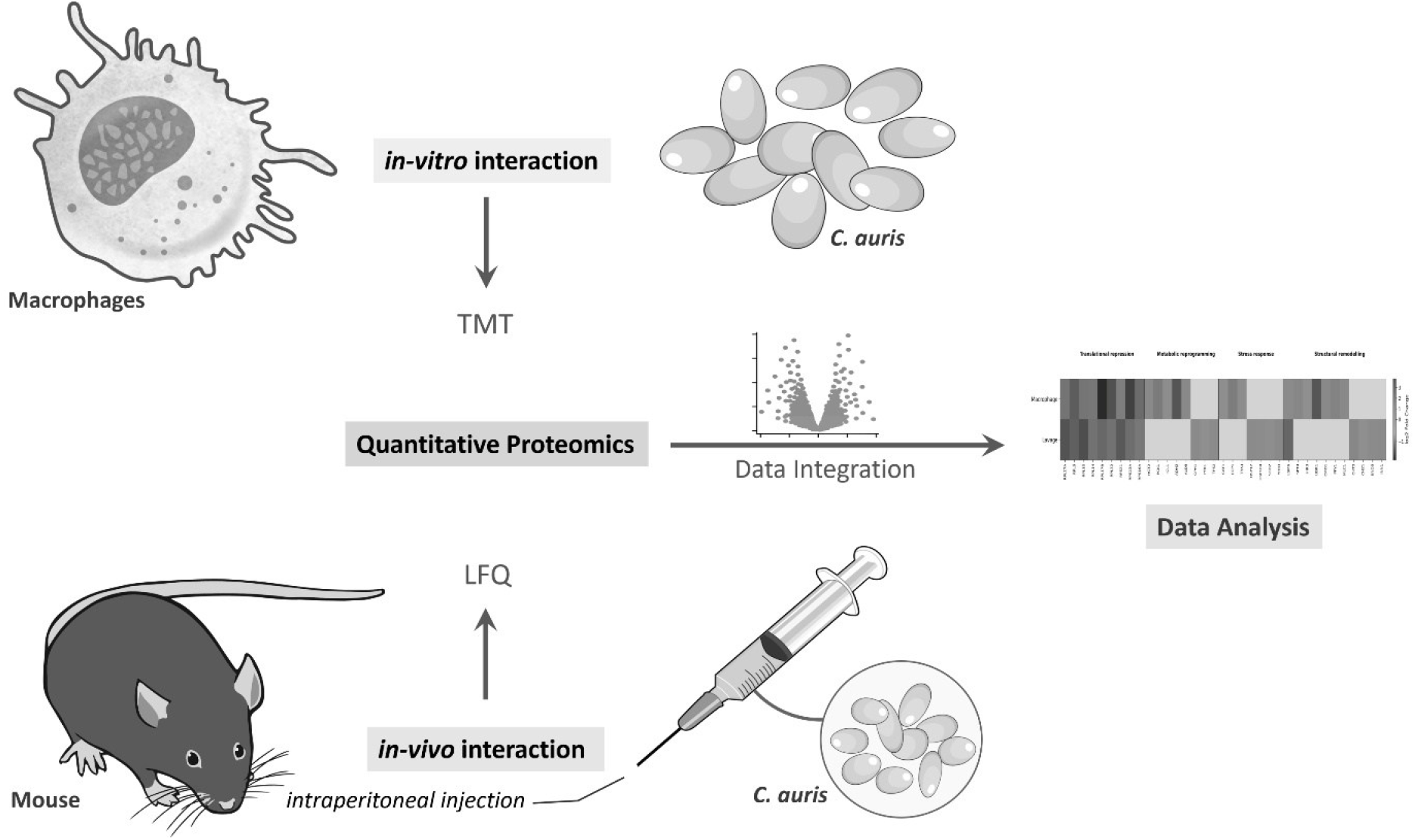
Experimental workflow applied to investigate *C. auris* host-pathogen interaction.

## 2 Materials and methods

### 2.1 Fungal culture

In this study we used *C. auris* isolate 1133/P/13-R (clade I), candida cells were routinely cultured in YPD liquid media (1% yeast extract, 2% peptone, and 2% glucose) at 30°C with constant shaking at 200 rpm.

### 2.2 Ethical statement

All animal experiments were conducted in accordance with approval from the Ethics Committee of the Medical University of Vienna and the Austrian Federal Ministry of Science and Research adhering to European legislation for animal experimentation [11].

### 2.3 Isolation of primary macrophages and *in-vitro* co-culture

Isolation and cultivation of primary bone marrow-derived macrophages (BMDM) was performed as described previously [11]. Briefly, after sacrificing 8-12 weeks old female C57BL/6 wild-type mice *(Mus musculus)* by cervical dislocation, the abdomen and the hind legs were sterilized with 70% ethanol. Followed by isolation of femurs and tibias, after which the bones were briefly rinsed with 70% ethanol and stored in 25mL phosphate-buffered saline (PBS) while moving onto another mouse. After all the bones were collected in PBS from all mice, they were aseptically separated at the knee joints. After this, the bones were transferred into sterile mortar and pestle with 10mL cold Dulbecco’s Modified Eagle Medium (DMEM) media for grinding. After through grinding, the entire suspension was passed through 0.22µM cell strainer. Cells were then harvested at 300g for 5 min at 4°C and incubated in red cell lysis buffer in the ratio of 500µL /leg for 2 min at room temperature. Afterwards, the reaction was stopped by the addition of 25mL cold DMEM media. Bone marrow cells were harvested at 300g for 5 min at 4 °C and resuspended L929-conditioned-BMDM media for differentiation and cell culture. For, co-culture assays, macrophages were counted by a CASY counter (Roche, Swiss) and seeded in 6-well plates with 1 ×10^5^ cells/well at multiplicity of infection (MOI= 10:1; Candida:Macrophage) and incubated in 2.5mL DMEM media at 37 °C, 5% CO_2_ incubator for 2h. Control cultures contained no immune cells. Experiments were performed as biological triplicates. Additionally, macrophage fungal kill efficiency was monitored at MOI = 1:1. Following 2h incubation, 500µL of 0.2% Triton X-100 was added into each well, after which, cells were gently scrapped from the wells using fresh cell scrappers and the suspensions were transferred into 15mL falcon tubes followed by 30 sec of vortex. An aliquot of this was utilized for plating in YPD agar plates for CFU determination with appropriate dilutions. The cell suspensions in 15mL falcon tubes were then centrifuged at 3000g for 5 min, followed by 3x wash in sterile ice-cold PBS, with the final cell suspension being processed for proteome isolation.

### 2.4 Mouse infection and re-isolation of fungal cells from lavage

For *in-vivo* experiments, 6-8 weeks old female C57BL/6 mice were injected intraperitoneally with 7.5 × 10^7^ fungal cells suspended in 300µL sterile PBS using 27Gauge needles. Control mice received 300µL sterile PBS alone. Experiments were performed as biological triplicates. At 2h post-infection, mice were sacrificed and peritoneal lavage was performed by washing the peritoneal cavity three times with sterile PBS. Lavage fluids were pooled to a final 30mL total volume per animal in sterile 50mL falcon tubes. An aliquot of this was utilized for plating in YPD plates for CFU determination with appropriate dilutions. After this, lavage fluids were centrifuged at 300g for 5 min to separate cellular material from supernatant. Following which, the supernatant fraction was then further centrifuged at 3000g for 10 min at 4°C to pellet fungal cells and any residual cellular material. After this the majority of the supernatant was carefully pipetted out using 10mL and 1mL tips, leaving behind approximately 200µL, undisturbed and retained just above the visible pellet at the bottom of the falcon tube. From this 200µL, an aliquot of 10µL was utilized for visual examination by light microscopy (20x-40x magnification) to confirm the presence of fungal cells. Following this, 300µL of candida lysis buffer was added to the 200µL pallet sample. After which, the total volume of 500µL sample containing re-isolated fungal cells were then processed for proteome isolation.

### 2.5 Proteome isolation

Following fungal interaction *in-vitro* and *in-vivo*, protein was isolated from candida cells. Briefly, candida cells were centrifuged at 3000g for 5 min, followed by ice cold PBS wash-3x after which the cell pallet was resuspended in 500μL of ice-cold candida lysis buffer (1% sodium deoxycholate (SDC), 100mM Tris-HCl, 150mM NaCl, 1mM PMSF, 1mM EDTA, 1x cOmplete™ Protease Inhibitor (Roche, Swiss)) in 1.5mL screw cap tubes (Sarstedt, Germany). Next, the candida cells were subjected to mechanical disruption by glass beads using FastPrep™-24 5G Bead Beating (Fischer Scientific, USA). Following which, a small hole was punctured at the bottom of the screw cap tube using red-hot flamed needle and the screw cap tube was then inserted into 1.5mL Eppendorf tube and centrifuge at 3000g for 2 min at 4°C to separate the lysate from the glass beads. The lysate collected in the Eppendorf tube was then subjected to acetone precipitation by adding four volumes of ice-cold 100% acetone followed by 2h incubation at -20°C. Post incubation, the samples was centrifuged at 3000g for 5min to obtain the resulting protein pallet and supernatant acetone was discarded. The protein pallet was air dried on ice and was further processed for mass spectrometric analysis.

### 2.6 Label free quantitative proteomics of *in-vivo* lavage samples

#### 2.6.1 Sample preparation for mass spectrometry analysis

The protein pellets were resuspended in 300µL 4%(w/v) SDS, 100mM Tris/HCl pH 8.5, and incubated at 95° C for 5 min. The lysate was clarified by centrifugation at 16000g for 10min at 30° C. The supernatant was transferred to a new tube and the protein concentration was measured using 600nm protein assay kit (Pierce) with SDS compatibility reagents. 50µg protein was reduced by adding 30µL of 1M dithiothreitol (DTT) and heated at 95° C for 5 min. Then it was diluted with 9x 8M urea in 100mM Tris/HCl pH 8.5, transferred to the FASP filter, and centrifuged for 20min at 12000g. Then it was washed with 200µL 8 M urea in 100 mM Tris/HCl pH 8.5 for 20min at 12000g. 100µL 50mM Iodacetamide in 100mM Tris/HCl pH 8.5 was added and shaken for 1min and incubated 30min in the dark at room temperature. It was centrifuged for 10min at 12000g followed by washing twice with 200µL 8M urea in 100mM Tris/HCl pH 8.5 and three times with 100µL 50 mM ammonium bicarbonate (ABC). The filter was transferred to a new collection tube and 40µL 50mM ABC containing 1µg trypsin platinum (Promega) was added and kept at 37° C overnight. The digested peptides were collected by centrifuging for 15min at 12000g. The filter was washed with 40µL 50mM ABC, centrifuged for 15min at 12000g, and pooled. The digested peptides were acidified with 10µL 10% TFA and the peptides were desalted using C18 Stagetips [12] and MCX 96 well palte (Waters).

#### 2.6.2 Liquid chromatography-mass spectrometry analysis

Peptides were separated on an Ultimate 3000 RSLC nano-flow chromatography system (Thermo-Fisher), using a pre-column for sample loading (Acclaim PepMap C18, 5mm × 300µm, 5μm, Thermo-Fisher) and a C18 analytical column (Acclaim PepMap C18, 50cm × 75µm, 2μm, Thermo-Fisher), applying a segmented linear gradient from 2% to 35% and finally 80% solvent B (80 % acetonitrile, 0.1 % formic acid; solvent A 0.1 % formic acid) at a flow rate of 230 nL/min over 120 min. Eluting peptides were analyzed on an Exploris 480 Orbitrap mass spectrometer (Thermo Fisher) coupled to the column with a FAIMS pro ion-source (Thermo-Fisher) using coated emitter tips (PepSep, MSWil) with the following settings: The mass spectrometer was operated in DDA mode with two FAIMS compensation voltages (CV) set to -45 or -60 and 1.5 s cycle time per CV. The survey scans were obtained in a mass range of 350-1500 m/z, at a resolution of 60k at 200 m/z, and a normalized AGC target at 100%. The most intense ions were selected with an isolation width of 1.2 m/z, fragmented in the HCD cell at 28% collision energy, and the spectra recorded for max. 100 ms at a normalized AGC target of 100% and a resolution of 15k. Peptides with a charge of +2 to +6 were included for fragmentation, the peptide match feature was set to preferred, the exclude isotope feature was enabled, and selected precursors were dynamically excluded from repeated sampling for 45 seconds.

#### 2.6.3 Data analysis

Raw data were split into single cv using FreeStyle 1.8 SP2 and processed using the MaxQuant software package (version 1.6.17.0, [13]) and the Uniprot *Candida auris* reference proteome (version 2021.03, www.uniprot.org), as well as a database of most common contaminants. The search was performed with full trypsin specificity and a maximum of two missed cleavages at a protein and peptide spectrum match false discovery rate of 1%. Carbamidomethylation of cysteine residues were set as fixed, oxidation of methionine and N-terminal acetylation as variable modifications. For label-free quantification the “match between runs” feature and the LFQ function were activated - all other parameters were left at default. MaxQuant output tables were further processed in R using Cassiopeia_LFQ (https://github.com/maxperutzlabs-ms/Cassiopeia_LFQ). Reverse database identifications, contaminant proteins, protein groups identified only by a modified peptide, protein groups with less than two quantitative values in one experimental group, and protein groups with less than 2 razor peptides were removed for further analysis. Missing values were replaced by randomly drawing data points from a normal distribution modeled on the whole dataset (data mean shifted by -1.8 standard deviations, width of distribution of 0.3 standard deviations). Differences between groups were statistically evaluated using the LIMMA package [14] at 5% FDR (Benjamini-Hochberg).

### 2.7 TMT label based quantitative proteomics of *in-vitro* fungal cells

#### 2.7.1 Sample preparation for mass spectrometry analysis

The protein pellets were resuspended in 100µL 8M urea 50mM ammonium bicarbonate, with sonication and vortex. The protein was reduced by adding 3.2µL of 250mM dithiothreitol (DTT) in 50mM ammonium bicarbonate and kept at 37° C for 30 min, followed by adding 3.2µL 500 mM Iodacetamide in 50mM ammonium bicarbonate and incubated 30min in the dark at room temperature. The alkylation reaction was quenched by adding 1.6µL 250 mM DTT for 10 min. The urea concentration was reduced to 4M using 50 mM ABC. Samples were pre-digested for 3 hours at 37°C with Lys-C (FUJIFILM Wako Pure Chemical Corporation, 125-02543) at an enzyme-to-substrate ratio of 1:50 and subsequently reduced to 1M Urea using 50mM ABC and digested overnight at 37°C with sequencing grade trypsin (Trypsin Gold, Mass Spec Grade, Promega, V5280) at an enzyme-to-substrate ratio of 1:50. Digests were stopped by acidification with TFA (Thermo Scientific, 28903) (0.5% final concentration) and desalted on a HLB 96-well plate (Waters). Peptide concentrations were determined and adjusted according to UV chromatogram peaks obtained with an UltiMate 3000 Dual LC nano-HPLC System (Dionex, Thermo Fisher Scientific), equipped with a monolithic C18 column (Phenomenex). Desalted samples were concentrated in a SpeedVac concentrator (Eppendorf) and subsequently lyophilized overnight. Lyophilized peptides were dissolved in 50μl 100mM TEAB (Sigma). 450μg of each TMT-10Plex reagent (Thermo Fisher Scientific) were dissolved in 30μl of 100% anhydrous acetonitrile and 30μl was added to the peptide/TEAB mixes at 1:4 sample:label ratio. Labels used: 126C: candia_1; 127N: coculture_1; 127C: candia_2; 128N: coculture_2; 128C: candia_3; 129N: coculture_3; 129C: empty; 130N:mouse_1; 130C:mouse_2; 130C:mouse_3;. Samples were labeled for 60 min at RT. 0.1μl of each sample were pooled, mixed with 10 μl 0.1% TFA, and analyzed by mass spectrometry (MS) to check labeling efficiency. The labeling was repeated twice to make sure all channels have a labeling efficiency > 98%. For quenching, 8μl of 5% hydroxylamine was added and the reaction was incubated for 25 min at RT. Samples were pooled and subsequently desalted with Sep-Pak tC-18 cartridges (Waters). Desalted samples were dried for 30 min in a SpeedVac vacuum centrifuge and subsequently lyophilized overnight. The sample was high pH fractionated using Pierce high pH reversed-phase peptide fractionation kit. Nine fractions were collected and lyophilized overnight.

#### 2.7.2 Liquid chromatography-mass spectrometry analysis

Peptides were separated on an Ultimate 3000 RSLC nano-flow chromatography system (Thermo-Fisher), using a pre-column for sample loading (Acclaim PepMap C18, 5 mm × 300 µm, 5 μm, Thermo-Fisher) and a C18 analytical column (Acclaim PepMap C18, 50 cm × 75 µm, 2 μm, Thermo-Fisher), applying a segmented linear gradient from 2% to 35% and finally 80% solvent B (80 % acetonitrile, 0.1 % formic acid; solvent A 0.1 % formic acid) at a flow rate of 230 nL/min over 180 min. Eluting peptides were analyzed on an Orbitrap Eclipse mass spectrometer (Thermo Fisher) coupled to the column with a FAIMS pro ion-source (Thermo-Fisher) using coated emitter tips (PepSep, MSWil) with the following settings: The mass spectrometer was operated in DDA mode with three FAIMS compensation voltages (CV) set to -40, -55 or -70 and 1.2 s cycle time per CV. The survey scans were obtained in a mass range of 375-1500 m/z, at a resolution of 120k at 200 m/z, and a normalized AGC target at 100%. The most intense ions were selected with an isolation width of 1.2 m/z, fragmented in the HCD cell at 30% collision energy, and the spectra recorded for max. 50 ms at a normalized AGC target of 100% in the iontrap. Peptides with a charge of +2 to +6 were included for fragmentation, the peptide match feature was set to preferred, the exclude isotope feature was enabled, and selected precursors were dynamically excluded from repeated sampling for 45 seconds. High-resolution MS1 survey scans were acquired in the Orbitrap analyzer. Precursors were selected for fragmentation by CID in the linear ion trap (MS2). Real-Time Search was enabled to perform on-the-fly database searching of MS2 spectra using a sample FASTA database, with fix modifications of carbamidomethyl(C), TMT10plex (Kn) and variable modification oxidation(M) and deamidation (NQ). Only MS2 spectra passing the peptide-level FDR threshold (≤5%) triggered an MS3 event, thereby eliminating TMT quantification of unidentified or co-isolated precursors. MS3 scans utilized 10 SPS notches to co-isolate fragment ions for HCD fragmentation and Orbitrap detection of TMT reporter ions (m/z 110–500).

#### 2.7.3 Data analysis

Raw data were split into single cv using FreeStyle 1.8 SP2 and processed using the MaxQuant software package (version 2.0.3.0, [13]) and the Uniprot *Candida auris* reference proteome (version 2021.03, www.uniprot.org), as well as a database of most common contaminants. The search was performed in MS3 TMT10 reporter ion mode with isobaric label correction, with full trypsin specificity and a maximum of two missed cleavages at a protein and peptide spectrum match false discovery rate of 1%. Carbamidomethylation of cysteine residues were set as fixed, oxidation of methionine and N-terminal acetylation as variable modifications. For label-free quantification the “match between runs” feature and the LFQ function were activated - all other parameters were left at default. MaxQuant output tables were further processed in R using Cassiopeia_LFQ (https://github.com/maxperutzlabs-ms/Cassiopeia_LFQ). Reverse database identifications, contaminant proteins, protein groups identified only by a modified peptide, protein groups with less than two quantitative values in one experimental group, and protein groups with less than 2 razor peptides were removed for further analysis. The protein intensities were median normalized using only candia protein intensities. Missing values were replaced by randomly drawing data points from a normal distribution modeled on the whole dataset (data mean shifted by -1.8 standard deviations, width of distribution of 0.3 standard deviations). Differences between groups were statistically evaluated using the LIMMA package [14] at 5% FDR (Benjamini-Hochberg).

### 2.8 Proteomics data deposition

The mass spectrometry proteomics data have been deposited to the ProteomeXchange Consortium via the PRIDE partner repository [15] with the dataset identifier PXD074745 for label free proteomics and PXD074762 for TMT labelled proteomics.

## 3. Results

### 3.1 Interaction of *C. auris* with macrophages *in-vitro* and *in-vivo* mouse

To determine the susceptibility of *C. auris* to immune-mediated killing during the initial stages of infection, we evaluated fungal kill-kinetics at 2h under both *in-vitro* and *in-vivo* interaction conditions. During interaction with macrophages *in-vitro* at MOI of 1, *C. auris* was susceptible to immune killing, reflecting the efficiency of macrophages (Figure 2B). However, with an MOI of 10 fungal cells, *C. auris* persisted with no significant killing by immune cells (Figure 2A). Indicating numerical fungal burden influences kill-kinetics of immune cells. In contrast, during, *in-vivo* condition, significant *C. auris* reduction was observed (Figure 2C), indicating a more potent immune mediated killing within the murine host.

**Figure 2.**
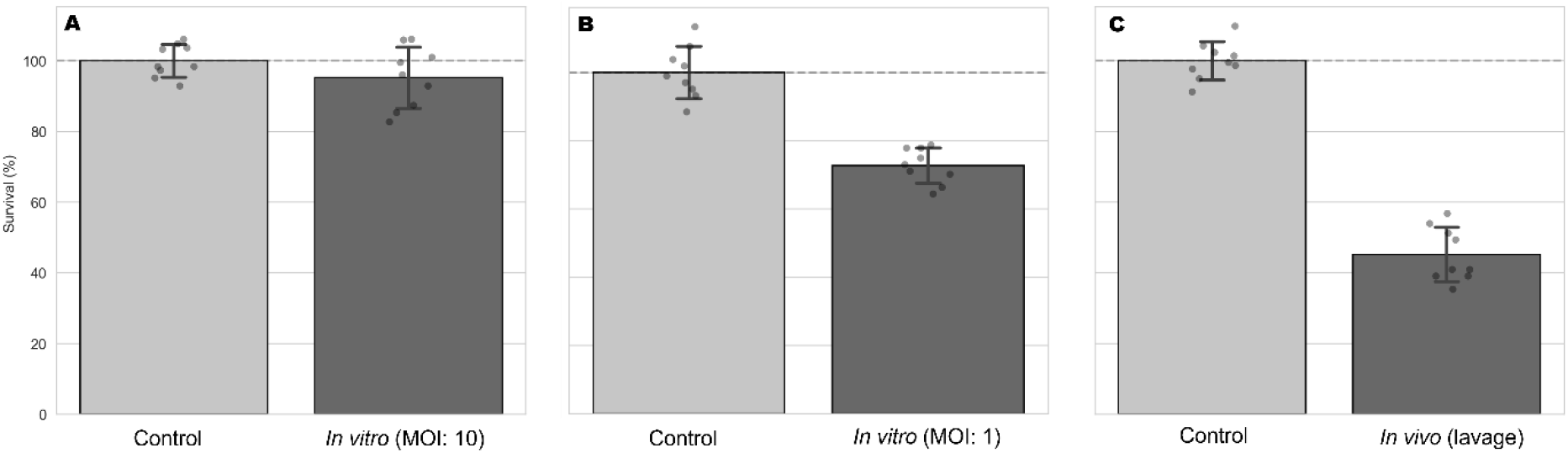
Early-phase kill-kinetics of *C. auris* were assessed during *in-vitro* and *in-vivo* conditions with 2h incubation. **(A)** *C. auris* survival during its interaction with macrophages at MOI 10:1 (candida:macrophage), no significant difference in survival was observed compared to the control (p = 0.449). **(B)** *C. auris* killing during its interaction with macrophages at MOI: 1, significant killing was noticed (p < 0.001). **(C)** *C. auris* survival *in-vivo*, significant killing of fungal cells compared to the control (p < 0.001). Statistical significance between the control and experimental groups was determined using an unpaired two-tailed T-test.

### 3.2 Proteome dynamics of *C. auris* during *in-vitro* and *in-vivo* interaction

To characterize the proteome landscape of *C. auris* during its interaction with the host, we applied a quantitative proteomics strategy on re-isolated fungal cells upon their interaction with macrophages *in-vitro* and *in-vivo* in mice. A total of 3034 fungal proteins were detected upon *in-vitro* interaction with macrophages (Supplementary Table 1). Of these, 1073 proteins were significantly differentially abundant, reflecting alterations in fungal proteome dynamics. Among the differentially abundant proteins (DAPs), 67 proteins were upregulated, while 82 proteins were downregulated (Figure 3A). The proteome landscape of *C. auris* during *in-vivo* interaction revealed a total of 2484 proteins, with 272 DAPs, where 53 proteins were upregulated and 135 proteins were downregulated (Figure 3B). Reflecting a similar shift in fungal proteome dynamics upon interaction with the host *in-vivo*.

**Figure 3.**
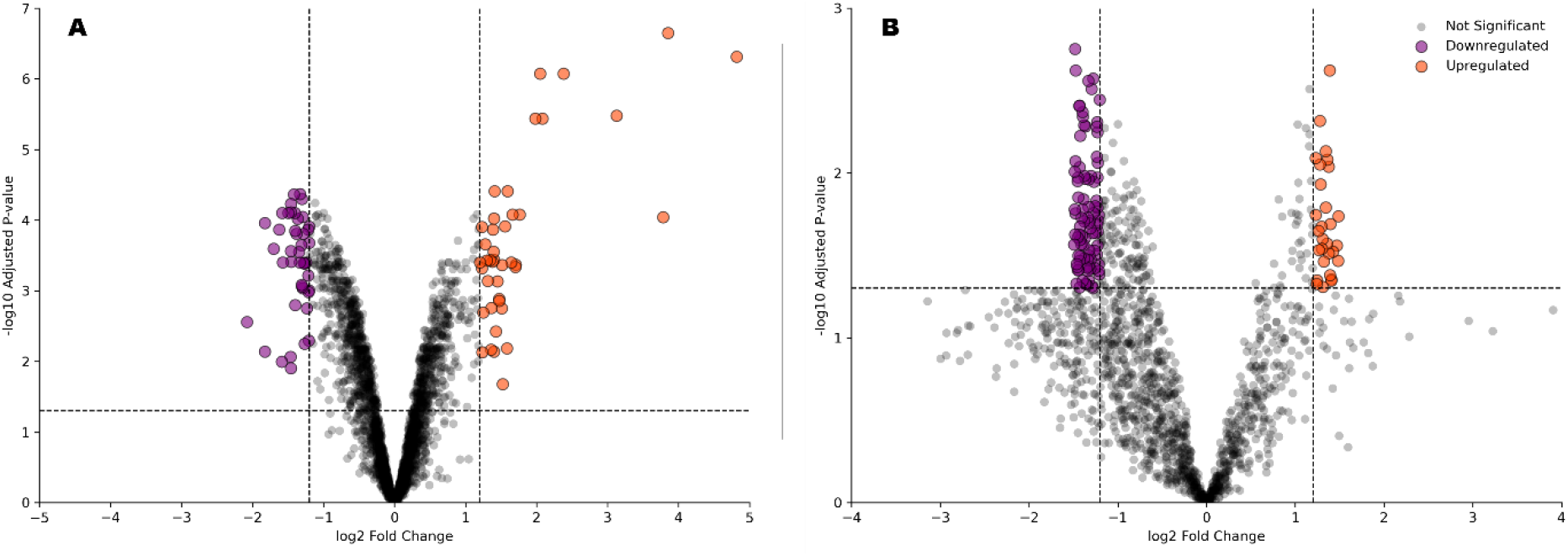
Volcano plots showing proteomic dynamics of *C. auris* during *in-vitro* macrophage and *in-vivo* lavage interactions. **(A)** Differentially abundant proteins (DAPs) during *C. auris* interaction with macrophages *in-vitro*. A total of 3034 fungal proteins were identified with 1073 significant DAPs where 67 proteins were upregulated and 82 were downregulated (adj. P.val ≤ 0.05, log FC ± 1). **(B)** DAPs of *C. auris* upon interaction *in-vivo* in mouse. A total of 2484 *C. auris* proteins were identified with 272 significant DAPs where 53 proteins were upregulated and 135 proteins were downregulated (adj. P.val ≤ 0.05, log FC ± 1).

### 3.3 Functional categorization of selected differentially abundant proteins

Selected proteins that were differentially abundant were assessed for their potential role in mediating early fungal-pathogen interactions. Functional categories associated with translational control, metabolic processes, stress response, and structural remodelling under both *in-vitro* and *in-vivo* conditions (Figure 4) were determined based on annotations from Candida Genome Database [16] and manual curations. Results indicated a common downregulation of multiple ribosomal proteins such as *Rpl27a, Rpl3, Rpl10, Rpl14, Rpl17b, Rpl32, Rps21, Rps23a*, and *Rps16a*. Proteins associated with metabolic adaptation were differentially abundant under both environments. During interaction with macrophages *in-vitro*, upregulated proteins included *Hgt2, Pck1, Icl1, Adh2*, and *Ald5*, whereas *in-vivo* lavage samples showed modulation of *Gph1, Pfk1*, and *Tps2*. These alterations suggest condition-specific adjustments to utilize carbon. Stress-associated proteins were also upregulated, including *Cat1, Ccp1, Trx1, Sod2, Yhb1, Hsp12*, and *Hsp104*, consistent with host-associated stress exposure. Finally, proteins linked to structural remodelling were upregulated including *Cmd1, Pfy1, Mlc1, End3, Inn1, Cht3, Lsm6, Spt4, Hir3*, and *Ubr1*, indicating proteomic alterations influencing cytoskeletal organization. Altogether, these data suggest a coordinated and context-dependent proteome modulation during *C. auris* host-pathogen interaction.

**Figure 4.**
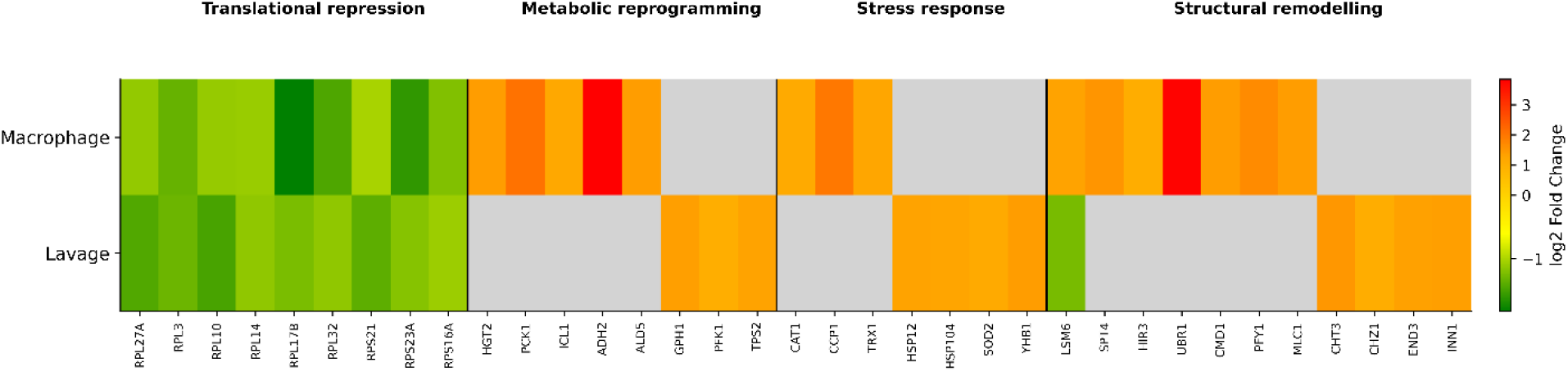
Functional categorization of differentially abundant proteins in macrophage *(in-vitro)* and lavage *(in-vivo)* interaction conditions. Heatmap showing log_2_ fold changes of selected proteins grouped by biological function: translational repression, metabolic reprogramming, stress response, and structural remodelling.

## 4 Discussion

The infection models used in this study demonstrate that immune-mediated killing of *C. auris* is influenced by both fungal burden and host context. During the *in-vitro* interaction assay, macrophages significantly reduced fungal viability at MOI of 1, however no significant killing was observed at MOI of 10, indicating increased fungal burden may limit effective immune clearance. The persistence of *C. auris* at higher fungal densities implies the existence of robust immune-evasion strategies, potentially driven by rapid molecular adaptions and collective microbial defence, with the latter reminiscent of biofilm like condition. In contrast, upon systemic *in-vivo* infection, a significant reduction in fungal viability was observed, suggestive of a larger context of co-ordinated immune response within the host. In order to further delve into the host-pathogen dynamics of *C. auris* we applied a quantitative proteomics methodology on re-isolated fungal cells from the *in-vitro* and *in-vivo* infection models, which reflected a broad proteome remodelling by *C. auris* in response to the host. Of note, it is important to point out the complexities of establishing experimental infection models and their integration thereof, as such the results of proteomics analysis should serve as elucidations warranting further functional validations.

## 4.1 Translational repression

A conserved feature of the *C. auris* proteome dynamics during *in-vitro* and *in-vivo* host interaction was a coordinated downregulation of proteins involved in translation and ribosome biogenesis. Multiple ribosomal proteins were downregulated, including *Rpl27a, Rpl3, Rpl10, Rpl14, Rpl17b, Rpl32, Rps21, Rps23a*, and *Rps16a* [16,17]. Such a broad repression of translational factors indicates a conserved shift toward a stress-adapted state. Translational attenuation is consistent with cellular stress responses and reflects adaptive strategy under hostile conditions [18,19]. The shared translational suppression across both *in-vitro* and *in-vivo* environments suggests that exposure to the host-immune pressure trigger a core adaptive response. Rather than promoting proliferation, *C. auris* appears to prioritize survival, metabolic efficiency, and stress tolerance.

### 4.2 Metabolic reprogramming

Exposure to host environment triggered a wide-spectrum metabolic rewiring by *C. auris*. During macrophage interaction *in-vitro*, enzymes such as high-affinity glucose transporter *Hgt2*, gluconeogenesis enzyme *Pck1* and glyoxylate cycle enzyme *Icl1* were upregulated, suggesting a shift towards a dynamic carbon utilization [20–22]. The enzymes alcohol dehydrogenase *Adh2* and aldehyde dehydrogenase *Ald5* were also upregulated suggesting the additional involvement of ethanol metabolism [23,24]. The activation of these pathways can be linked to nutrient limitation and intracellular survival and is therefore, indicative of metabolic reprogramming upon macrophage internalization, warranting *C. auris* to rapidly utilize of a variety of metabolic substrates. During *in-vivo* interaction, enzymes such as *Gph1, Pfk1, Tps2* were upregulated. The enzyme *Gph1* is essential for glycogen synthesis in *C. albicans* and plays a role in virulence [25], the pathogen can accumulate carbohydrates in the form of glycogen and mobilize these intracellular reserves when external nutrient availability becomes limited. The upregulation of *Gph1 in-vivo* suggests that *C. auris* may also engage in similar adaptive mechanism, whereby host-imposed nutrient constraints may promote glycogen accumulation under nutrient-limited conditions. The enzyme *Pfk1* is a key regulator of glycolysis and has been reported to be a crucial virulence factor in *C. albicans* [26]. The modulation of *Pfk1 in-vivo* reflects potential adaptive response by *C. auris* for energy metabolism in response to host-imposed environmental constraints. Finally, the upregulation of *Tps2*, a key enzyme in the trehalose biosynthetic pathway further supports the notion of adaptive metabolic remodelling by *C. auris* under host-associated stress [27]. Recent work has demonstrated the role of *Tps2* in *C. auris*, including its role in regulating stress tolerance, antifungal susceptibility, and virulence [27]. The enrichment of a wide range of metabolic proteins associated with gluconeogenesis, glyoxylate cycle and trehalose metabolism therefore suggests an integrated and rapid metabolic adaptation by *C. auris* upon immediate exposure to host-stress. Such a pattern is consistent with metabolic flexibility displayed by numerous pathogens including *Candida* species in a host environment where nutrients may be limited.

### 4.3 Stress response

Host-immune responses create hostile microenvironments to fend-off invading pathogens, in particular immune cells can generate high levels of reactive oxygen/nitrogen species (ROS/RNS) to impose collective stress [28]. Therefore, in order to establish a successful infection, pathogens such as *C. auris* must counter such stress responses. In agreement with this, under both *in-vitro* and *in-vivo* conditions an immediate activation of a robust stress response was observed. During macrophage interaction *in-vitro*, oxidative stress regulators including *Cat1, Ccp1*, and *Trx1* were upregulated; enzymes known to mediate ROS detoxification in *C. albicans* [29–33]. In contrast, *in-vivo* exposure induced a broader stress profile marked by *Hsp12, Hsp104, Sod2*, and the nitric oxide scavenger *Yhb1*. The proteins *Hsp12* and *Hsp104* has been reported to chaperone virulence and survival in *C. albicans* during host interaction, particularly to mitigate osmotic and thermal stress [34], while *Sod2* mitigates ROS toxicity [33]. The upregulation of *Yhb1* suggests adaptation to host-derived nitric oxide, paralleling nitrosative stress responses described in *C. albicans* host-pathogen interaction [35]. Collectively, induction of these stress mediators underscores a coordinated oxidative and general stress adaptation strategy, enabling *C. auris* to withstand immediate ROS/RNS exposure within hostile host microenvironments.

### 4.4 Structural remodelling

Morphological adaptation emerged as another distinguishing feature during *C. auris* interaction with the host *in-vitro* and *in-vivo*. During *in-vitro* interaction several molecules mediating structural response were upregulated such as *Lsm6, Spt4, Hir3, Ubr1, Cmd1, Pfy1* and *Mlc1*. The proteins associated with *Lsm6* and *Spt4* were reported to play role for filamentation in *C. albicans* [36]. The *Hir3* is a component of the HIR complex, which in *C. albicans* has been implicated in regulating morphological plasticity, from yeast-to-hyphae, a key virulence-associated trait. [37]. Additional regulators such as *Ubr1* and *Cmd1* encoding for calmodulin have also been reported to mediate hyphal development in *C. albicans* [38,39]. The upregulation of these yeast-to-hyphae associated factors in our dataset suggests the induction of pseudohyphal growth in *C. auris* immediately upon macrophage interaction. Such morphological transitions have been previously described in *C. auris* under genotoxic stress [40], raising the possibility that immune-mediated stress may induce fungal morphogenesis. Such a structural plasticity was further accentuated by the upregulation of myosin component *Mlc1* and profilin *Pfy1*, these were reported to be crucial molecules modulating fungal growth, cell wall integrity and virulence in *C. albicans* [41,42]. Altogether, these findings indicate coordinated cytoskeletal and morphogenetic remodelling of *C. auris* during macrophage interaction. During *in-vivo* interaction proteins such as *Cht3, Chz1, End3* and *Inn1* were upregulated. The enzyme *Cht3* is a key chitinase involved in cell wall remodelling and contributes to virulence in *C. albicans* [43,44]. The *Chz1* is an essential chitin synthase associated protein, which has been shown regulate for cell wall integrity [45], whereas *Inn1* has been reported to regulate chitin synthase [46]. As such, both of these proteins can modulate cell morphogenesis by regulating chitin turnover in *C. auirs*. Additionally, such chitin associated molecules have been reported to be downregulated following antifungal exposure [47], suggesting their potential role against stressors. Therefore, the induction of chitin-remodelling proteins *in-vivo* likely contributes to adaptive cell wall remodelling in *C. auris* in response to host-imposed stress.

### 4.5 Potential moonlighting function

Moonlighting proteins perform distinct, often unrelated functions depending on cellular localization or microenvironmental context, and are increasingly recognized as important virulence factors in fungal pathogens [48,49]. Several proteins identified in this study may possess additional ‘moonlighting’ function. For example, the glycolytic enzyme *Pgk1* have been detected on the cell surface where it can bind plasminogen [50], suggesting functions beyond central carbon metabolism. Similarly, pyruvate decarboxylase *Pdc11* has been reported to localize on cellular surface in *C. albicans*, indicating potential non-metabolic roles [51]. The protein *Gna1* essential for UDP-N-acetylglucosamine synthesis have also been identified on cell surface plus reported to mediate fungal survival in host [51,52]. Likewise, *Yke2* has been associated with both carbon metabolism and morphogenesis, thereby priming metabolic function to structural remodelling [53]. Furthermore, *Trx1* has been known for its multifaceted role in promoting fungal survival in the host, in addition to mitigating oxidative stress, [29]. The upregulation of such proteins by *C. auris* may therefore reflect not only metabolic engagement but also context-dependent roles that drive host-pathogen interactions.

## 5 Conclusion

Collectively our findings suggest, that immediate exposure to host environments drives a coordinated and multifaceted adaptive response in *C. auris*. A conserved translational repression across both *in-vitro* and *in-vivo* conditions indicates a potential core adaptive response perhaps prioritizing survival over proliferation. Apart from core response, context dependent divergence in fungal stress and metabolic response was observed. Interaction with macrophages *in-vitro* was characterized by metabolic reprogramming toward alternative carbon utilization and cytoskeletal remodelling consistent with intracellular adaptation, whereas *in-vivo* interaction leaned towards carbohydrate metabolism and a co-ordinated cell wall remodelling. At the same time, robust oxidative and nitrosative stress responses underscore the capacity of *C. auris* to withstand host-imposed ROS/RNS pressure. Structural remodelling through morphogenetic regulation and chitin-associated enzymes further highlights adaptive plasticity of *C. auris* during host-pathogen interaction. The identification of moonlighting proteins also suggests that metabolic and stress-associated molecules may additionally contribute to host-pathogen interaction beyond their conventional roles. Altogether, our findings indicate that early adaptation of *C. auris* to host environments is accompanied by coordinated modulation of translational machinery, metabolic rewiring, stress-response, and structural remodelling, a framework warranting further investigation.

## 6 Acknowledgement

This study was funded by the Austrian Science Fund (FWF) Principal Investigator Projects: CandidOmics-P33425 (Grant DOI: 10.55776/P33425) awarded to Dr. Rounik Mazumdar. Proteomics analyses were performed by the Mass Spectrometry Facility at Max Perutz Labs using the VBCF instrument pool. We would like to thank Karl Kuchler and lab members for their support. For the purpose of open access, the authors have applied a CC BY public copyright license.

## 7 CRediT author statement

Conceptualization: RM. Methodology: RM. Software: RM. Validation: RM, AB. Formal analysis: RM. Investigation: RM, AB. Resources: RM. Data curation: RM. Visualization: RM. Supervision: RM. Project administration: RM. Funding acquisition: RM. Writing-original draft: RM. Writing-Review & Editing: RM, AB. Both authors have approved the final manuscript.

## References

1. Bhargava A, Klamer K, Sharma M, Ortiz D, Saravolatz L. Candida auris: A Continuing Threat. Microorganisms. 2025;13:652. doi:10.3390/microorganisms13030652

2. Hayes JF. Candida auris: Epidemiology Update and a Review of Strategies to Prevent Spread. J Clin Med. 2024;13:6675. doi:10.3390/jcm13226675

3. The Lancet Microbe. Candida auris: new clade, same challenges. Lancet Microbe. 2024;5:100977. doi:10.1016/j.lanmic.2024.100977

4. De Gaetano S, Midiri A, Mancuso G, Avola MG, Biondo C. Candida auris Outbreaks: Current Status and Future Perspectives. Microorganisms. 2024;12:927. doi:10.3390/microorganisms12050927

5. Nett JE. Candida auris: An emerging pathogen “incognito”? Hogan DA, editor. PLoS Pathog. 2019;15:e1007638. doi:10.1371/journal.ppat.1007638

6. Sears D, Schwartz BS. Candida auris: An emerging multidrug-resistant pathogen. International Journal of Infectious Diseases. 2017;63:95–98. doi:10.1016/j.ijid.2017.08.017

7. Lockhart SR, Etienne KA, Vallabhaneni S, Farooqi J, Chowdhary A, Govender NP, et al. Simultaneous Emergence of Multidrug-Resistant Candida auris on 3 Continents Confirmed by Whole-Genome Sequencing and Epidemiological Analyses. Clinical Infectious Diseases. 2017;64:134–140. doi:10.1093/cid/ciw691

8. Mazumdar R, Bjelanovic A. A targeted drug-repurposing strategy identifies Tavaborole (Kerydin) as a potent fungistatic agent against Candida auris. 2026. doi:10.64898/2026.02.27.708474

9. Zhang Y, Zeng L, Huang X, Wang Y, Chen G, Moses M, et al. Targeting epigenetic regulators to overcome drug resistance in the emerging human fungal pathogen Candida auris. Nat Commun. 2025;16:4668. doi:10.1038/s41467-025-59898-6

10. Cha H, Won D, Bahn Y-S. Signaling pathways governing the pathobiological features and antifungal drug resistance of Candida auris. mBio. 2025;16. doi:10.1128/mbio.02475-23

11. Penninger P, Riedelberger M, Tsymala I, Arzani H, Jenull S, Kuchler K. Quantification of zinc intoxication of Candida glabrata after phagocytosis by primary macrophages. STAR Protoc. 2021;2:100352. doi:10.1016/j.xpro.2021.100352

12. Rappsilber J, Mann M, Ishihama Y. Protocol for micro-purification, enrichment, pre-fractionation and storage of peptides for proteomics using StageTips. Nat Protoc. 2007;2:1896–1906. doi:10.1038/nprot.2007.261

13. Tyanova S, Temu T, Cox J. The MaxQuant computational platform for mass spectrometry-based shotgun proteomics. Nat Protoc. 2016;11:2301–2319. doi:10.1038/nprot.2016.136

14. Ritchie ME, Phipson B, Wu D, Hu Y, Law CW, Shi W, et al. limma powers differential expression analyses for RNA-sequencing and microarray studies. Nucleic Acids Res. 2015;43:e47–e47. doi:10.1093/nar/gkv007

15. Perez-Riverol Y, Bai J, Bandla C, García-Seisdedos D, Hewapathirana S, Kamatchinathan S, et al. The PRIDE database resources in 2022: a hub for mass spectrometry-based proteomics evidences. Nucleic Acids Res. 2022;50:D543–D552. doi:10.1093/nar/gkab1038

16. Lew-Smith J, Binkley J, Sherlock G. The Candida Genome Database: annotation and visualization updates. Genetics. 2025;229. doi:10.1093/genetics/iyaf001

17. Shamsuzzaman M, Rahman N, Gregory B, Bommakanti A, Zengel JM, Bruno VM, et al. Inhibition of Ribosome Assembly and Ribosome Translation Has Distinctly Different Effects on Abundance and Paralogue Composition of Ribosomal Protein mRNAs in Saccharomyces cerevisiae. mSystems. 2023;8. doi:10.1128/msystems.01098-22

18. Flint A, Butcher J, Stintzi A. Stress Responses, Adaptation, and Virulence of Bacterial Pathogens During Host Gastrointestinal Colonization. Microbiol Spectr. 2016;4. doi:10.1128/microbiolspec.VMBF-0007-2015

19. Jobava R, Mao Y, Guan B-J, Hu D, Krokowski D, Chen C-W, et al. Adaptive translational pausing is a hallmark of the cellular response to severe environmental stress. Mol Cell. 2021;81:4191-4208.e8. doi:10.1016/j.molcel.2021.09.029

20. Chew SY, Chee WJY, Than LTL. The glyoxylate cycle and alternative carbon metabolism as metabolic adaptation strategies of Candida glabrata: perspectives from Candida albicans and Saccharomyces cerevisiae. J Biomed Sci. 2019;26:52. doi:10.1186/s12929-019-0546-5

21. Martin R, Albrecht-Eckardt D, Brunke S, Hube B, Hünniger K, Kurzai O. A Core Filamentation Response Network in Candida albicans Is Restricted to Eight Genes. PLoS One. 2013;8:e58613. doi:10.1371/journal.pone.0058613

22. Jakab Á, Balla N, Ragyák Á, Nagy F, Kovács F, Sajtos Z, et al. Transcriptional Profiling of the Candida auris Response to Exogenous Farnesol Exposure. mSphere. 2021;6. doi:10.1128/mSphere.00710-21

23. Pateman JAJ, Doy CH, Olsen JE, Norris U, Creaser EH, Hynes M. Regulation of alcohol dehydrogenase (ADH) and aldehyde dehydrogenase (AldDH) in Aspergillus nidulans. Proc R Soc Lond B Biol Sci. 1983;217:243–264. doi:10.1098/rspb.1983.0009

24. Gutiérrez-Corona JF, González-Hernández GA, Padilla-Guerrero IE, Olmedo-Monfil V, Martínez-Rocha AL, Patiño-Medina JA, et al. Fungal Alcohol Dehydrogenases: Physiological Function, Molecular Properties, Regulation of Their Production, and Biotechnological Potential. Cells. 2023;12:2239. doi:10.3390/cells12182239

25. Miao J, Regan J, Cai C, Palmer GE, Williams DL, Kruppa MD, et al. Glycogen Metabolism in Candida albicans Impacts Fitness and Virulence during Vulvovaginal and Invasive Candidiasis. mBio. 2023;14. doi:10.1128/mbio.00046-23

26. Luo J, Zhang Y, Zhang Y, Li S, Zhang H. Preliminary study of the lethal fungus Candida albicans phosphofructokinase-1 as a potential therapeutic target. Int J Biol Macromol. 2025;330:148039. doi:10.1016/j.ijbiomac.2025.148039

27. Zhu Q, Van de Velde S, Wijnants S, Carolus H, Jacobs S, Sofras D, et al. Accumulation of Trehalose 6-Phosphate in Candidozyma auris results in Decreased Echinocandin Resistance and Tolerance. Nat Commun. 2025;17:311. doi:10.1038/s41467-025-67022-x

28. Hitzler SUJ, Fernández-Fernández C, Montaño DE, Dietschmann A, Gresnigt MS. Microbial adaptive pathogenicity strategies to the host inflammatory environment. FEMS Microbiol Rev. 2025;49. doi:10.1093/femsre/fuae032

29. da Silva Dantas A, Patterson MJ, Smith DA, MacCallum DM, Erwig LP, Morgan BA, et al. Thioredoxin Regulates Multiple Hydrogen Peroxide-Induced Signaling Pathways in Candida albicans. Mol Cell Biol. 2010;30:4550–4563. doi:10.1128/MCB.00313-10

30. Enjalbert B, Smith DA, Cornell MJ, Alam I, Nicholls S, Brown AJP, et al. Role of the Hog1 Stress-activated Protein Kinase in the Global Transcriptional Response to Stress in the Fungal Pathogen Candida albicans. Mol Biol Cell. 2006;17:1018–1032. doi:10.1091/mbc.e05-06-0501

31. Wysong DR, Christin L, Sugar AM, Robbins PW, Diamond RD. Cloning and Sequencing of a Candida albicans Catalase Gene and Effects of Disruption of This Gene. Infect Immun. 1998;66:1953–1961. doi:10.1128/IAI.66.5.1953-1961.1998

32. Shin Y, Lee S, Ku M, Kwak M-K, Kang S-O. Cytochrome c peroxidase regulates intracellular reactive oxygen species and methylglyoxal via enzyme activities of erythroascorbate peroxidase and glutathione-related enzymes in Candida albicans. Int J Biochem Cell Biol. 2017;92:183–201. doi:10.1016/j.biocel.2017.10.004

33. Dantas A, Day A, Ikeh M, Kos I, Achan B, Quinn J. Oxidative Stress Responses in the Human Fungal Pathogen, Candida albicans. Biomolecules. 2015;5:142–165. doi:10.3390/biom5010142

34. Gong Y, Li T, Yu C, Sun S. Candida albicans Heat Shock Proteins and Hsps-Associated Signaling Pathways as Potential Antifungal Targets. Front Cell Infect Microbiol. 2017;7. doi:10.3389/fcimb.2017.00520

35. Ullmann BD, Myers H, Chiranand W, Lazzell AL, Zhao Q, Vega LA, et al. Inducible Defense Mechanism against Nitric Oxide in Candida albicans. Eukaryot Cell. 2004;3:715–723. doi:10.1128/EC.3.3.715-723.2004

36. Lash E, Maufrais C, Janbon G, Robbins N, Herzel L, Cowen LE. The spliceosome impacts morphogenesis in the human fungal pathogen Candida albicans. mBio. 2024;15. doi:10.1128/mbio.01535-24

37. Jenull S, Tscherner M, Gulati M, Nobile CJ, Chauhan N, Kuchler K. The Candida albicans HIR histone chaperone regulates the yeast-to-hyphae transition by controlling the sensitivity to morphogenesis signals. Sci Rep. 2017;7:8308. doi:10.1038/s41598-017-08239-9

38. Lu Y, Su C, Unoje O, Liu H. Quorum sensing controls hyphal initiation in Candida albicans through Ubr1-mediated protein degradation. Proceedings of the National Academy of Sciences. 2014;111:1975–1980. doi:10.1073/pnas.1318690111

39. Saporito SM, Sypherd PS. The isolation and characterization of a calmodulin-encoding gene (CMD1) from the dimorphic fungus Candida albicans. Gene. 1991;106:43–49. doi:10.1016/0378-1119(91)90564-R

40. Bravo Ruiz G, Ross ZK, Gow NAR, Lorenz A. Pseudohyphal Growth of the Emerging Pathogen Candida auris Is Triggered by Genotoxic Stress through the S Phase Checkpoint. mSphere. 2020;5. doi:10.1128/mSphere.00151-20

41. Sun X, Wang Y, Yang X, Xiang X, Zou L, Liu X, et al. Profilin Pfy1 is critical for cell wall integrity and virulence in Candida albicans. Microbiol Spectr. 2025;13. doi:10.1128/spectrum.02593-24

42. Puerner C, Serrano A, Wakade RS, Bassilana M, Arkowitz RA. A Myosin Light Chain Is Critical for Fungal Growth Robustness in Candida albicans. mBio. 2021;12. doi:10.1128/mBio.02528-21

43. Blázquez-Muñoz MT, Costa-Barbosa A, Alvarado M, Mendonça A, Benkhellat S, Vilanova M, et al. Loss of CHT3 in Candida albicans wild-type strains increases surface-exposed chitin and affects host-pathogen interaction. Front Cell Infect Microbiol. 2025;15. doi:10.3389/fcimb.2025.1654710

44. Dünkler A, Walther A, Specht CA, Wendland J. Candida albicans CHT3 encodes the functional homolog of the Cts1 chitinase of Saccharomyces cerevisiae. Fungal Genetics and Biology. 2005;42:935–947. doi:10.1016/j.fgb.2005.08.001

45. Munro CA, Winter K, Buchan A, Henry K, Becker JM, Brown AJP, et al. Chs1 of Candida albicans is an essential chitin synthase required for synthesis of the septum and for cell integrity. Mol Microbiol. 2001;39:1414–1426. doi:10.1046/j.1365-2958.2001.02347.x

46. Devrekanli A, Foltman M, Roncero C, Sanchez-Diaz A, Labib K. Inn1 and Cyk3 regulate chitin synthase during cytokinesis in budding yeasts. J Cell Sci. 2012. doi:10.1242/jcs.109157

47. Kaneko Y, Ohno H, Kohno S, Miyazaki Y. Micafungin Alters the Expression of Genes Related to Cell Wall Integrity in &lt;i&gt;Candida albicans&lt;/i&gt; Biofilms. Jpn J Infect Dis. 2010;63:63.355. doi:10.7883/yoken.63.355

48. Satala D, Karkowska-Kuleta J, Zelazna A, Rapala-Kozik M, Kozik A. Moonlighting Proteins at the Candidal Cell Surface. Microorganisms. 2020;8:1046. doi:10.3390/microorganisms8071046

49. Karkowska-Kuleta J, Satala D, Bochenska O, Rapala-Kozik M, Kozik A. Moonlighting proteins are variably exposed at the cell surfaces of Candida glabrata, Candida parapsilosis and Candida tropicalis under certain growth conditions. BMC Microbiol. 2019;19:149. doi:10.1186/s12866-019-1524-5

50. Chen S-M, Zou Z, Guo S-Y, Hou W-T, Qiu X-R, Zhang Y, et al. Preventing Candida albicans from subverting host plasminogen for invasive infection treatment. Emerg Microbes Infect. 2020;9:2417–2432. doi:10.1080/22221751.2020.1840927

51. Gil-Bona A, Parra-Giraldo CM, Hernáez ML, Reales-Calderon JA, Solis N V., Filler SG, et al. Candida albicans cell shaving uncovers new proteins involved in cell wall integrity, yeast to hypha transition, stress response and host–pathogen interaction. J Proteomics. 2015;127:340–351. doi:10.1016/j.jprot.2015.06.006

52. Mio T, Kokado M, Arisawa M, Yamada-Okabe H. Reduced virulence of Candida albicans mutants lacking the GNA1 gene encoding glucosamine-6-phosphate acetyltransferase. Microbiology (N Y). 2000;146:1753–1758. doi:10.1099/00221287-146-7-1753

53. Liboro K, Yu S-R, Lim J, So Y-S, Bahn Y-S, Eoh H, et al. Transcriptomic and Metabolomic Analysis Revealed Roles of Yck2 in Carbon Metabolism and Morphogenesis of Candida albicans. Front Cell Infect Microbiol. 2021;11. doi:10.3389/fcimb.2021.636834

